# Unbiased functional genetic screens reveal essential RNA modifications in human cancer and drug resistance

**DOI:** 10.1101/2024.07.13.603368

**Authors:** Cornelius Pauli, Michael Kienhöfer, Maximilian Felix Blank, Oguzhan Begik, Christian Rohde, Daniel Heid, Fu Xu, Sarah Miriam Naomi Zimmermann, Katharina Weidenauer, Sylvain Delaunay, Nadja Krall, Katrin Trunk, Duoduo Zhao, Fengbiao Zhou, Anke Heit-Mondrzyk, Uwe Platzbecker, Claudia Baldus, Hubert Serve, Martin Bornhäuser, Cathrine Broberg Vågbø, Salvador Aznar Benitah, Jeroen Krijgsveld, Eva Maria Novoa, Carsten Müller-Tidow, Michaela Frye

## Abstract

RNA modification pathways are mis-regulated in multiple types of human cancer. To comprehensively identify cancer-relevant RNA modifications and their regulators, we screened all 150 annotated human RNA modifying proteins across 18 different normal and cancer cell lines using a CRISPR-based genetic knockout system. Fifty RNA modifying proteins were essential for survival of at least one cell type. A third of these essential genes were amplified in 38 different human primary cancer types and potentially drive cancer growth. Unexpectedly, the number of essential genes per cell line varied considerably, and this variation was not due to tissue of origin. Instead, we found that cancer cell-specific mitochondrial metabolic plasticity was responsible for the unique requirement of certain RNA modifications. For example, leukemia cells with high intrinsic drug tolerance required mitochondrial flexibility to survive treatment with the anti-leukemic drugs cytarabine and venetoclax. Synthetic lethality screens revealed that drug-resistance is abolished by deleting the mitochondrial methyltransferase TRMT5, which is responsible for the formation of N1-methylguanosine (m^1^G) in the tRNA anticodon loop. In summary, our study identifies cancer-relevant RNA modifying enzymes, and reveals a novel promising drug target for therapy-resistant acute myeloid leukemia.

## Introduction

Many of the 170 known nucleotide modifications occurring in coding and non-coding RNAs orchestrate gene expression programs to regulate fundamental cellular functions ^1,2^. Most types of RNAs will carry biochemical modifications at some point in their lifetime to alter electrostatic charge, base pairing, secondary structure, and RNA-protein interactions ^3^. RNA modifications regulate transcriptional and post-transcriptional processes including RNA processing and stability, nuclear export, cellular localization and mRNA translation ^4^. Thus, RNA modifications serve as a molecular bridge between gene transcription and transcriptional output.

How RNA modifications regulate cellular functions is best-studied during acute stress responses induced by oxidative stress, DNA damage or anti-cancer drugs ^2,5–7^. In response to external stress stimuli, RNA modifying proteins cause degradation or stabilization of their target RNA, and thereby re-wire cellular proteomes even before transcriptional changes occur ^5–9^. Thus, RNA modifications sense alterations in the environmental and initiate cellular stress responses accordingly.

The ability to adapt to a changing micro-environment is crucial for cell fate decisions during development ^10^. Aberrant deposition of RNA modifications is frequently linked to human diseases such as neurological deficits and metabolic disorders including obesity and diabetes ^2,5–7^. Similarly, cancer cells continually and dynamically adapt to often deleterious micro-environments such as hypoxia or drugs treatments ^11^. RNA modifications affect tumorigenesis by altering gene expression and cellular processes. But in contrast to transcription factor networks that maintain cell identity, RNA modification pathways mediate cellular functions by modulating temporal and spatial gene expression programs. Aberrant deposition of RNA modifications disrupts stable gene expression programs and alters basic cellular functions including cell growth, survival, differentiation, and migration ^9^. Most aspects of tumorigenesis including initiation, progression, metastasis and drug resistance require distinct RNA modifying events to ensure survival and growth ^5,9,12^.

Thus, RNA modifications are important regulators of cancer development and progression, and targeting them can provide new therapeutic strategies for cancer treatment. Yet, the majority of current efforts are limited to N6-methyladenosine (m^6^A) modifying enzymes. Here, we set out to identify all cancer-relevant RNA modifications in an unbiased manner through CRISPR/Cas9-based genetic screening of all currently known 150 RNA modifying proteins ^13^. We identify at least 15 essential RNA modifying proteins showing genomic amplifications across 38 different tumour types. Many of these proteins have not been implicated in cancer before. Although we find no evidence for essential tissue-specific RNA modifications, we reveal that distinct nucleotide modifications are required to fuel cell-specific metabolic demands. We show that mitochondria require TRMT5-mediated m^1^G modifications in tRNA anticodon loops to efficiently adapt to high energy demands required for survival in response to cancer drug treatment. In summary, chemical nucleotide modifications in RNA are rich targets for the development of novel therapeutic strategies.

## Results

### Target discovery screens identify essential RNA modifications in human cancer cells

To identify cancer-relevant RNA modifying proteins (RMPs) in an unbiased manner, we generated pooled guide RNA (gRNA) libraries targeting all currently known RMPs (n=150) ^13^ and eight control gRNAs targeting known oncogenes and essential genes. Each gene was targeted by 8 single guide (sg) RNAs. In addition, we included 250 small nucleolar (sno) RNAs, and 50 non-targeting sgRNAs as controls (Figure 1A; Supplementary Table 1).

**Figure 1.**
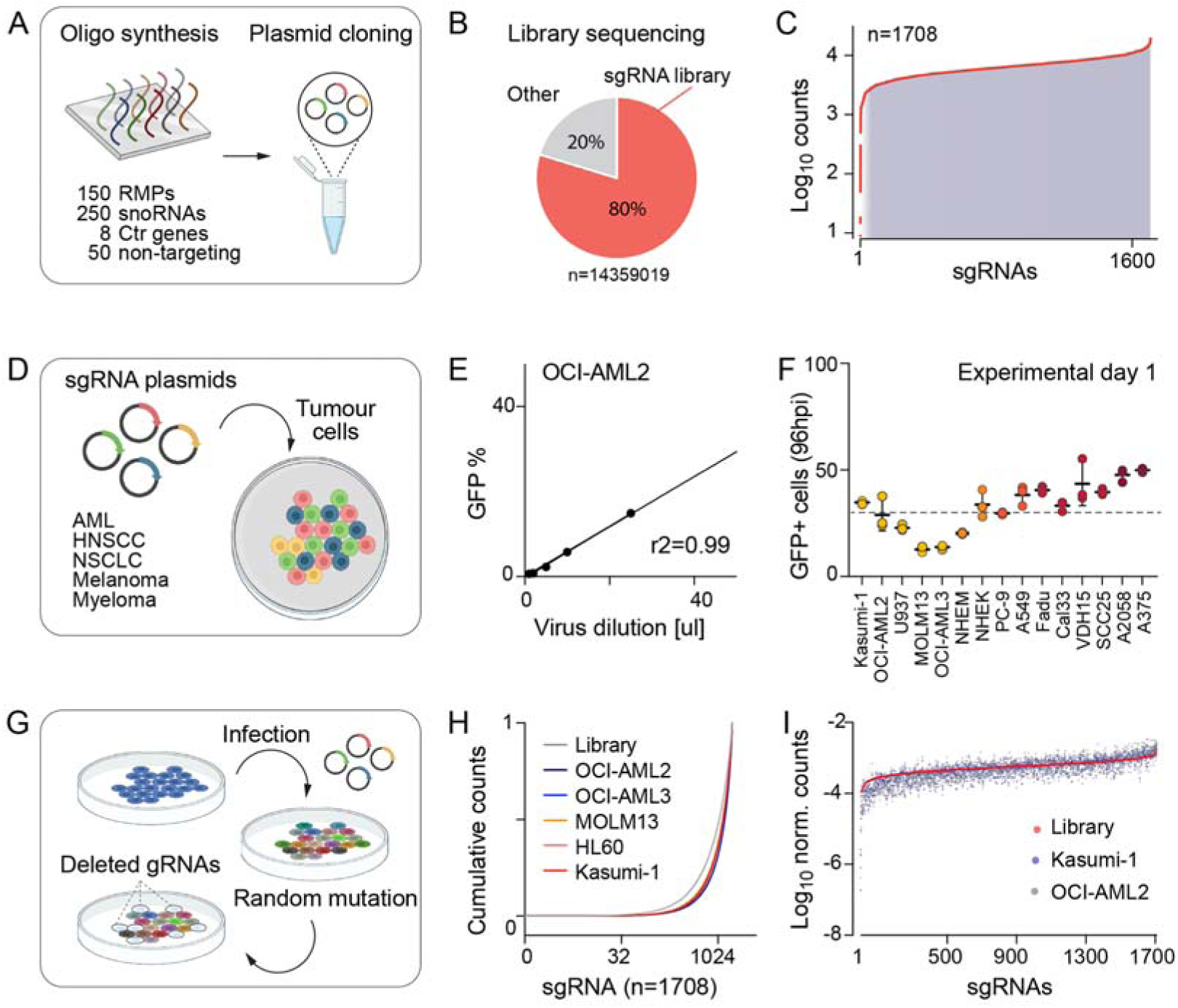
CRISPR Cas9 screen targeting human RNA modifying proteins. (**A**) Illustration of pooled guide RNA (gRNA) library targeting 150 RNA modifying proteins (RMPs) (n=150 covered by 8 single guide RNAs (sgRNA) each), small nucleolar RNAs (snoRNAs) (n=250 covered by 1-2 sgRNAs each), 8 control genes (Ctr) (n=8 covered by 8 single guide RNAs each) and 50 non-targeting gRNAs serving as negative control. (**B, C**) Library plasmid sequence read distribution (B) and coverage (C) across all 1708 sgRNAs. (**D**) Illustration of infected cancer cell lines. AML: acute myeloid leukemia, HNSCC: head and neck squamous cell carcinoma, NSCLC: non-small-cell lung carcinoma. (**E, F**) Representative virus titration in OCI-AML2 cells to a MOI of 0.3 (E) and representative infection rates (GFP-positive cells) at experimental day one 96 hours after infection (F). (**G**) Illustration of dropout screen performed in 18 cell lines. (**H, I**) Representative cumulative gRNA sequencing read counts in the indicated cell lines (H) and normalized Log_10_ read counts across all 1708 sgRNAs (I) 22 days after infections. Mean ± SD (F).

Sequencing of the library confirmed a high on-target rate and equal read count distribution across all sgRNAs (Figure 1B, C). We infected the library into 16 different cancer cell lines including acute myeloid leukemia (AML), head and neck squamous cell carcinoma (HNSCC), non-small-cell lung carcinoma (NSCLC), melanoma, and multiple myeloma. In addition, we screened primary healthy melanocytes (NHEM) and keratinocytes (NHEK) (Figure 1D; Supplementary Table 2). For each cell line, the virus was titrated to a MOI of 0.3 leading to an infection rate of around 30% of cells at day one of the experiment (Figure 1E, F; Supplementary Figure 1A-K). Then, we performed dropout viability screens to identify essential RMPs (Figure 1G). After eight and 22 days, we quantified enrichment or depletion of the gRNAs relative to day one of the experiment using high throughput sequencing. A minimum coverage of 500 x was maintained throughout all screening experiments (Figure 1H, I; Supplementary Figure 1L).

About one third of gRNAs targeting RMPs were consistently reduced across all 18 cell lines after 22 days of culture (Figure 2A). Similar to the oncogene MYC, TRMT112 was strictly required for cellular survival (Figure 2A, B). TRMT112 activates both rRNA and tRNA methyltransferases. While the METTL5 – TRMT112 complex forms m^6^A in 18S rRNA, the THUMPD3 - TRMT112 complex installs m^2^G into cytoplasmic tRNAs ^14,15^. In contrast, only a small number of gRNAs were enriched indicating a growth advantage when the targeted RMP was depleted (Figure 2A). Control gRNAs targeting the tumour suppressor TP53 were the most enhanced as expected (Figure 2A, B). We obtained a similar dependency on RNA modifying proteins after eight days of culture (Supplementary Figure 2A, B). Thus, our targeted CRISPR Cas9-based screen identified 50 essential RNA modifying proteins across 18 different cell lines.

**Figure 2.**
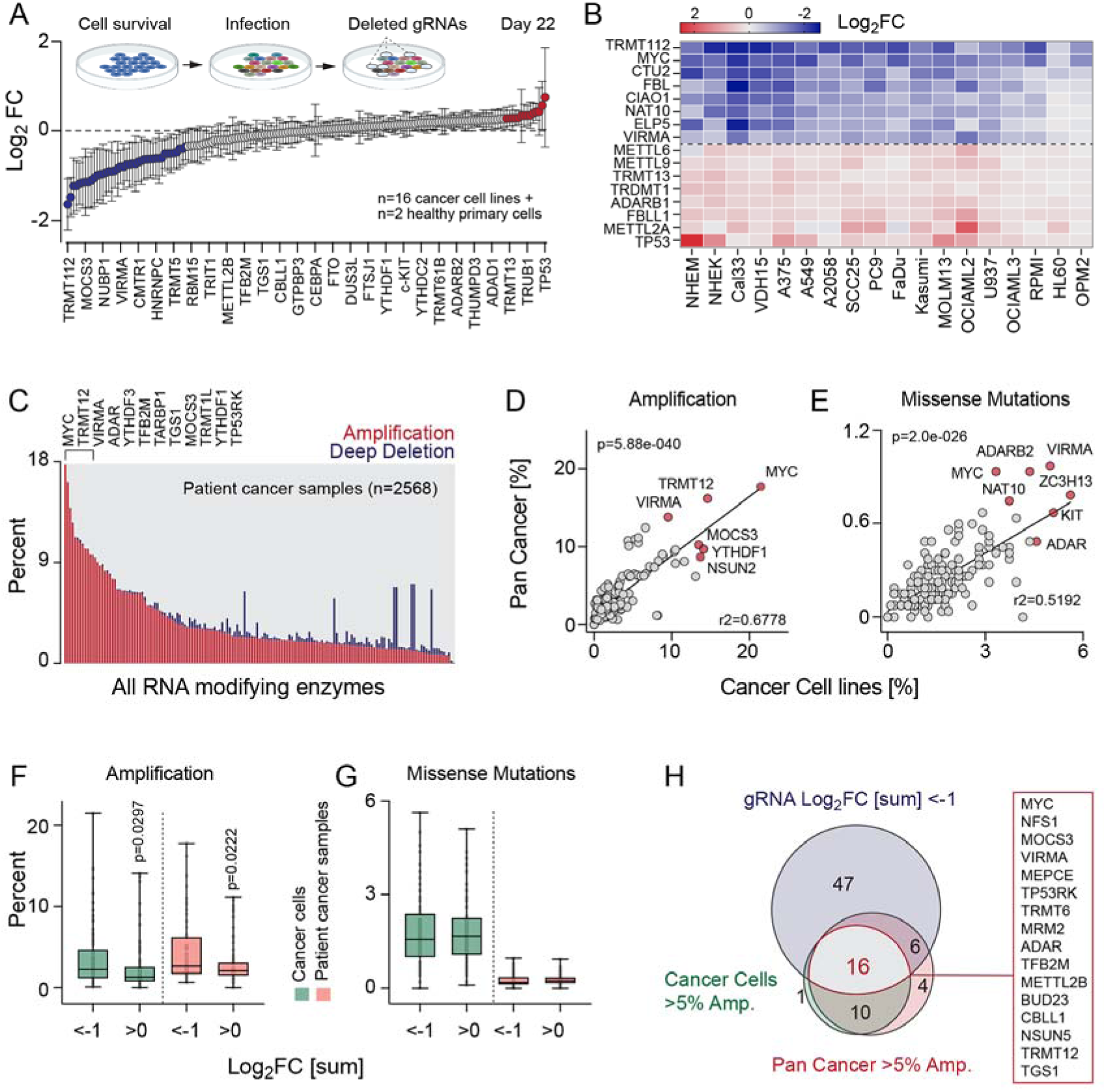
CRISPR/Cas9 screen identifies pan cancer-relevant RNA modifying proteins. **(A)** Scheme of dropout screen (top) and average log_2_ fold-change (FC) of 150 human RNA modifying proteins (RMPs) including positive controls in 18 cell lines (bottom) at day 22 of the experiment. Shown is every sixth RMP. (**B**) Heatmap of the top and bottom eight RMPs shown in (A). (**C**) Percentage of tumour samples showing genetic amplification (red) or deep deletions (blue) of 150 RMPs including positive controls. RMPs were covered in 2568 samples spanning 38 different tumour types (cBioPortal). Top 12 amplified RMPs are indicated (top). (**D, E**) Correlation of percentage of genetic amplification (D) or missense mutations (E) in 150 RMP genes including positive controls in patient cancer samples (n=2568) and cancer cell lines (n=961). Top six genes are highlighted in red. (**F, G**) Percentage of amplification (D) or missense mutations (E) in 150 RMP genes grouped into RMPs showing log_2_FC sum <−1 or >0 in patient cancer samples (red) and cancer cell lines (green). (**H**) Overlap of RMPs (red box) showing log_2_FC sum <−1 (n=69) and genetically amplified in > 5% patient cancer samples (red) and cancer cell lines (green). Mean ± SD (A). Box plots show minimum, first quartile, median, third quartile, and maximum (F,G). Two-tailed unpaired t-test (F).

### Pan cancer-relevant RNA modifying proteins

To identify pan cancer-relevant RNA modifications, we first identified all RMPs showing alterations in cancer. We used two datasets, the pan cancer study of whole genomes covering 38 tumour types and the Cancer Cell Line Encyclopedia (Figure 2C; Supplementary Figure 2C-E) ^16,17^. Five RMPs (TRMT12, VIRMA, MOCS3, YTHDF1, NSUN2) were genetically amplified in more than 10% of patient cancer samples and cell lines (Figure 2C,D; Supplementary Figure 2C). An extensive copy number gain of NSUN2 had already been reported in breast cancer ^18^. In contrast, deep deletions were inconsistent between cell lines and patient cancer samples and therefore, might represent cell culture artefacts (Figure 2C; Supplementary Figure 2C; blue bars). In addition, at least five RMPs were consistently mutated (predicted passenger mutations) in cancer (Figure 2E; Supplementary Figure 2D,E).

To test whether amplified or mutated RMPs were also required for cancer cell survival, we separated the RMPs into two groups: negatively (Log_2_FC sum < −1) or positively (Log_2_FC sum > 0) enriched in the dropout screens. Essential RMPs were indeed more likely to be genetically amplified in cancer (Figure 2F), while frequently mutated RMPs were not prone to be required for cancer cell survival (Figure 2G).

In summary, we identified at least 15 essential RMPs that were also amplified in more than 5% of all cancer patient samples across 38 tumour types (Figure 2H).

### RNA-modifying proteins function independently of tissue of origin

To test for essential cell type-specific RMPs, we clustered the 18 cell lines according to their depleted gRNAs (Figure 3A; Supplementary Figure 3A). In this dataset, 54 out of 161 depleted proteins were essential in at least one tested cell line (Figure 3B). While normal primary cells (NHEK, NHEM) separated from cancer cells, the first principal component was not driven by tissue of origin but instead, reflected the total number of essential RMPs per cell line ranging from 4 to 38 (Figure 3A-D). Since our data indicated that RMPs function largely independent of tissue of origin, we asked whether any RMPs were tissue-specifically expressed in healthy cells. Less than half of all RMPs showed some tissue-specificity according to single cell RNA sequencing data obtained from 25 human tissues (Human Protein Atlas) (Figure 3E; left panel; Supplementary Table 3). Since we also found that essential RMPs were slightly enriched for proteins with low tissue-specificity (Figure 3E, middle and right panel), we asked whether the cancer lines expressed only subsets of RMPs.

**Figure 3.**
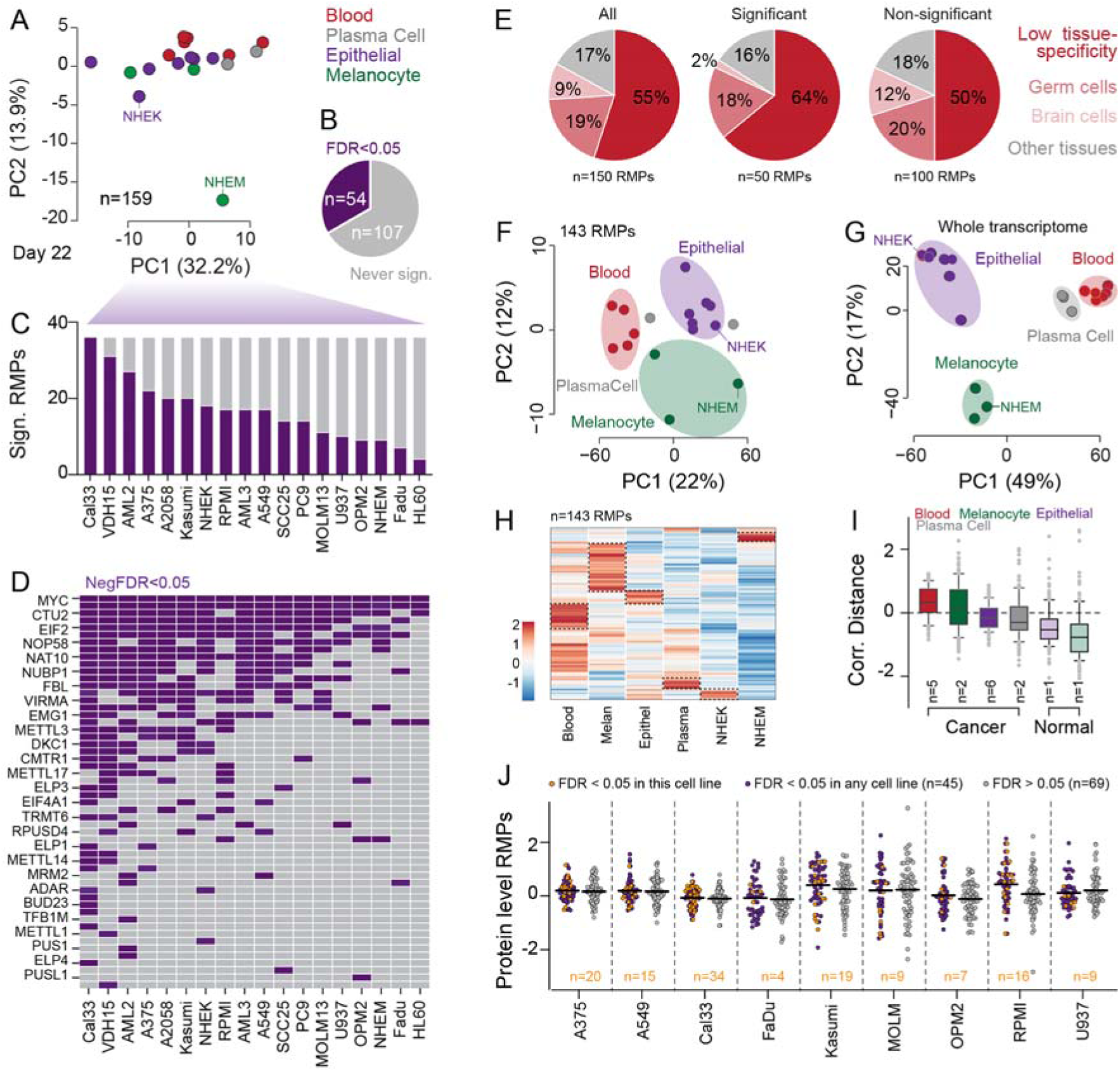
RNA modifying proteins function are largely independent of tissue of origin. **(A)** Principal component analysis (PCA) using log_2_ fold change (FC) of 159 gRNAs in 18 cell lines originating from blood, plasma cells, epithelial cells or skin melanocytes at day 22 of the experiment. (**B**) Number of significantly (FDR<0.5; purple) depleted gRNAs in at least one cell line. (**C, D**) Total number (C) and list of RNA modifying proteins (RMPs) (D) corresponding to significantly (FDR<0.05) depleted gRNAs in the indicated cell lines. (**E**) Proportion of all (left) (n=150), essential (middle) (n=50) or non-essential (right) (n=100) RMPs showing weak or strong tissue-specific expression (Human Protein Atlas). (**F,G**) PCA using RMP expression (n=143) (F) or whole transcriptomes (G) (normalized log_2_ FPKM) in 18 cell lines (n=3 sequencing reactions per cell line). (**H**) Heatmap of RMP-expression shown in (F) average over tissue or showing primary normal cells (NHEK, NHEM). (**I**) Averaged correlation distance of RMP-expression shown in (H) in cancer and normal cells. (**J**) Protein level of RMPs in the indicated cell lines grouped by gRNAs significantly (FDR<0.05) depleted in the indicated cell line (purple) or in any tested cell line (orange) or never significantly (FDR>0.05) depleted in any cell line (grey). Shown are 114 out of 159 proteins present in all cell lines. Box plots show minimum, first quartile, median, third quartile, and maximum (I).

Transcriptional profiling clearly separated the 17 cell lines by tissue of origin, and the vast majority of RMPs (>95%) were expressed across all cell lines (Figure 3F, G; Supplementary Figure 3B-E). Each tissue preferentially expressed distinct sets of RMPs (Figure 3H; Supplementary Figure 3F, G). Nevertheless, expression of all RMPs was on average higher in cancer lines when compared to their normal counterparts (Figure 3I), and blood cells expressed the highest levels of RMPs (Figure 3I; Supplementary Figure 3F). However, RMP-expression levels were poor indicators for essential cellular functions because RNA and protein levels were both uncorrelated to abundance of gRNAs in dropout screens (Figure 3J; Supplementary Figure 3H-M).

In summary, we showed that expression of RMPs was on average higher in cancer than normal cells, and we successfully identified pan cancer-relevant RMPs.

### Synthetic lethality screen identifies essential roles of TRMT5 in drug tolerance

Our data so far revealed that the number of essential RMPs per cell line varied substantially even within one cancer type such as acute myeloid leukaemia (AML) (Figure 4A). RNA modifications are often linked to cellular stresses such as oxidative stress, hypoxia, DNA damage or drug treatments. Therefore, we considered the possibility that the high heterogeneity between cancer lines reflected cell-specific adaptations to disease state or treatment condition. To test for inherent stress tolerances, we treated AML lines with cytarabine (AraC), a commonly used chemotherapy drug. We found that intrinsic drug tolerance varied widely, and OCI-AML2/3 cells exhibited an over 100-fold greater tolerance towards AraC than all other tested lines (Figure 4B). Drug tolerance indeed correlated with similar sets of essential RMPs (Figure 4C; Supplementary Table 4).

**Figure 4.**
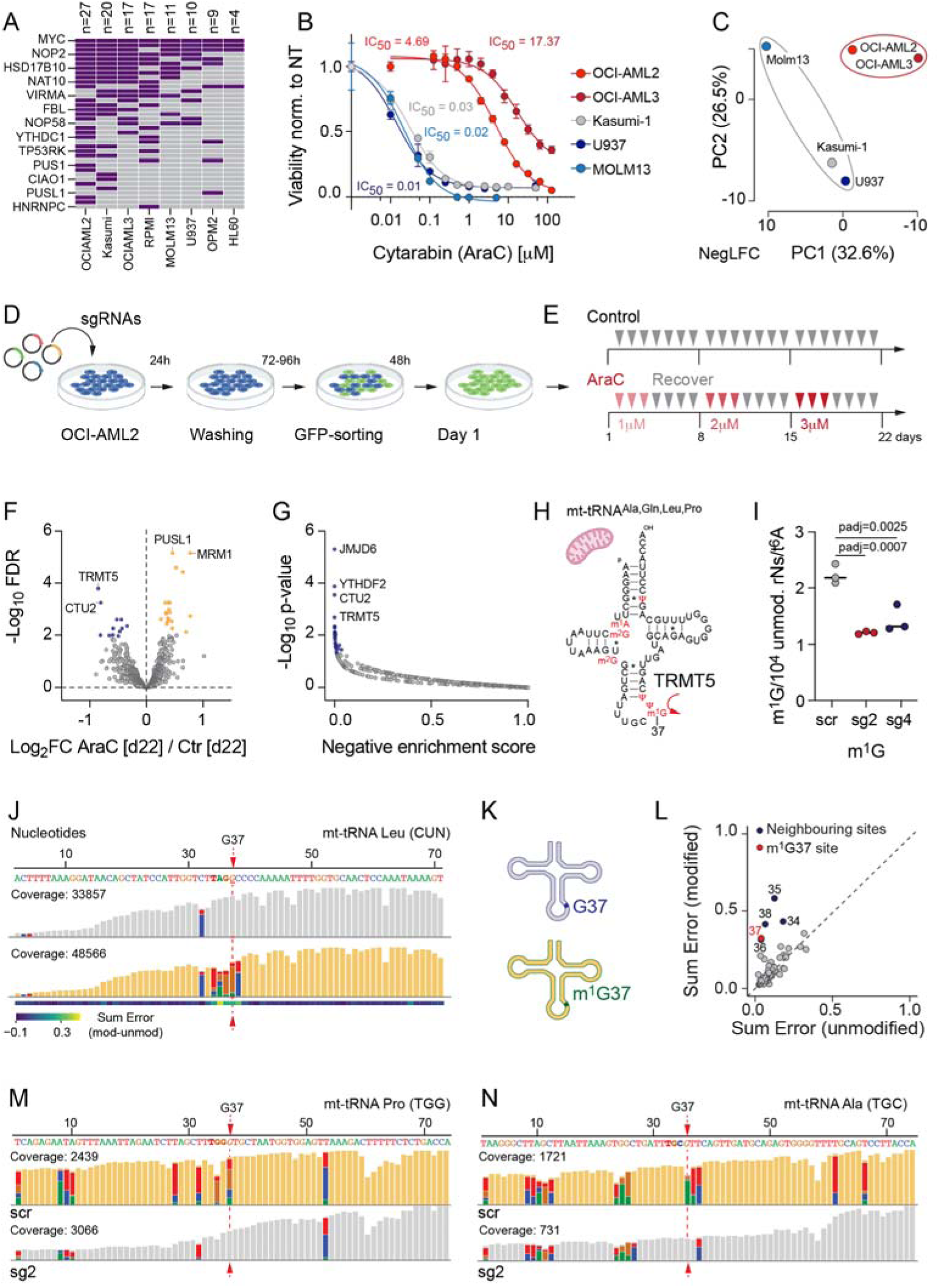
Synthetic lethality screen identifies novel vulnerabilities in drug resistance. **(A)** Heatmap of significantly (FDR<0.05) depleted gRNAs in blood cancer cell lines. (**B,C**) Dose-response curve examining the effect of different concentrations of cytarabine (AraC) in blood cancer lines with corresponding IC50 values (B) and cluster analyses using negative log_2_ fold changes of 159 gRNAs in 8 cancer lines originating from blood (C). PC: Principal component. (**D,E**) Illustration of synthetic lethality screen (D) and treatment regimen with AraC (E) in OCI-AML2 cells. (**F,G**) Volcano plot showing statistical significance (FDR) versus log_2_ fold change (FC) of sgRNA abundance (F) and negative enrichment score of averaged gRNAs per gene (G) identifying essential RMPs in AraC-treated cells after 22 days in culture. (**H,I**) Illustration of mitochondrial (mt) tRNAs methylated by TRMT5 at position G37 (H) and quantification of m^1^G levels in mitochondrial tRNAs using mass spectrometry (I). Modification levels are normalized to N6-threonyl-carbamoylation (t^6^A), a universal modification of adenosine 37 in tRNAs. (**J,K**) IGV snapshot (J) and illustration (K) of synthetic unmodified (top; grey) and m^1^G-modified (bottom; orange) mt-RNA ^Leu(CNN)^. (**L**) Comparison of summed error frequencies for each base of modified and unmodified synthetic mt-RNA ^Leu(CNN)^. (**M,N**) IGV snapshot of mt-tRNA^Pro(TGG)^ (M) and mt-tRNA^Ala(TGC)^ (N) from OCI-AML2 control (scr; top) and TRMT5-knockout (sg2) cells. Nucleotides with mismatch frequencies >0.05 are coloured (J,M,N).

To identify RMPs required for resistance towards AraC, we performed a synthetic lethality screen by repeatedly exposing OCI-AML2 cells to an increasing concentration of AraC (Figure 4D, E). To exclude RMPs commonly required for survival and proliferation, we compared treated versus untreated cells both cultured for 22 days (Figure 4F; Supplementary Figure 4A). Single gRNAs (sgRNA) targeting TRMT5 and CTU2 respectively were the most significantly depleted (Figure 4F; Supplementary Figure 4B,C). Similarly, also the averaged negative enrichment score of all eight sgRNAs targeting TRMT5 and CTU2 respectively were significantly reduced (Figure 4G). Both RMPs modify tRNA anti-codon loops. Human TRMT5 methylates the N1 position of guanosine 37 (G37) in mitochondrial tRNAs (Figure 4H), while CTU2 is responsible for the 2-thiolation of cytosolic tRNAs ^19–21^. Since AML cells are highly dependent on mitochondrial oxidative phosphorylation (OXPHOS) for energy production ^22^, we next asked how TRMT5 was linked to cellular survival and drug resistance.

### TRMT5 mediates formation of m^1^G in mitochondrial and cytosolic tRNAs

We selected two sgRNAs (sg2 and sg4) and generated TRMT5-knockout cell lines for the drug-tolerant OCI-AML2 and -sensitive MOLM13 lines (Supplementary Figure 4D,E). We isolated mitochondrial tRNAs and confirmed a two-fold reduction of m^1^G levels in the knockout cells by mass spectrometry (Figure 4H,I). Residual m^1^G levels were likely due to TRMT10A-mediated methylation at the G9 position in tRNAs.

To quantify all m^1^G37 levels in their sequence-specific contexts, we performed nanopore direct RNA sequencing (DRS). As proof-of-principle, we first sequenced modified and unmodified synthetic mt-tRNA^Leu^, a known target of TRMT5 (Figure 4J-L; Supplementary Table 5), showing that m^1^G modifications are efficiently detected using DRS. Then, we compared the mt-tRNA modification profile of TRMT5-knockout to control cells. We measured a decrease of m^1^G37 in all four known human mt-tRNA targets (Figure 4M,N; Supplementary Figure 4F; Supplementary Table 6). In addition, the majority of cytosolic G37-containing tRNAs showed a reduction in m^1^G signals in the absence of TRMT5 (Supplementary Figure 4G). We conclude that human TRMT5 methylated both mitochondrial and nuclear-encoded tRNAs at position G37.

### TRMT5 is required for mRNA translation of OXPHOS components

Mitochondrial functions and high oxidative phosphorylation (OXPHOS) status are hallmarks of AraC-resistant persisting leukemic cells ^23^. Therefore, we speculated that the enhanced drug tolerance of OCI-AML2 cells may directly be linked to altered mitochondrial metabolic activity. Up-regulation of OXPHOS requires increased translation of mitochondrial-encoded genes of the electron transport chain. Modifications in the mt-tRNA anticodon loop including m^1^G ensure efficient codon-specific translation ^24^.

To test whether TRMT5 was required for mRNA translation, we measured OP-puromycin incorporation into mitochondrial elongating polypeptide chains using flow cytometry (Figure 5A). Depletion of TRMT5 significantly reduced mitochondrial protein synthesis (Figure 5B). Accordingly, basal and maximal oxygen consumption rates were also decreased in TRMT5 knockout cells (Figure 5C, D). Total proteome analyses of TRMT5-knockout cell lines confirmed reduction of electron transport chain proteins and revealed increased levels of proteins involved in the amino acid starvation stress response (Supplementary Figure 5A,B; Supplementary Table 7). In conclusion, TRMT5 was required for optimal use of OXPHOS for energy production.

**Figure 5.**
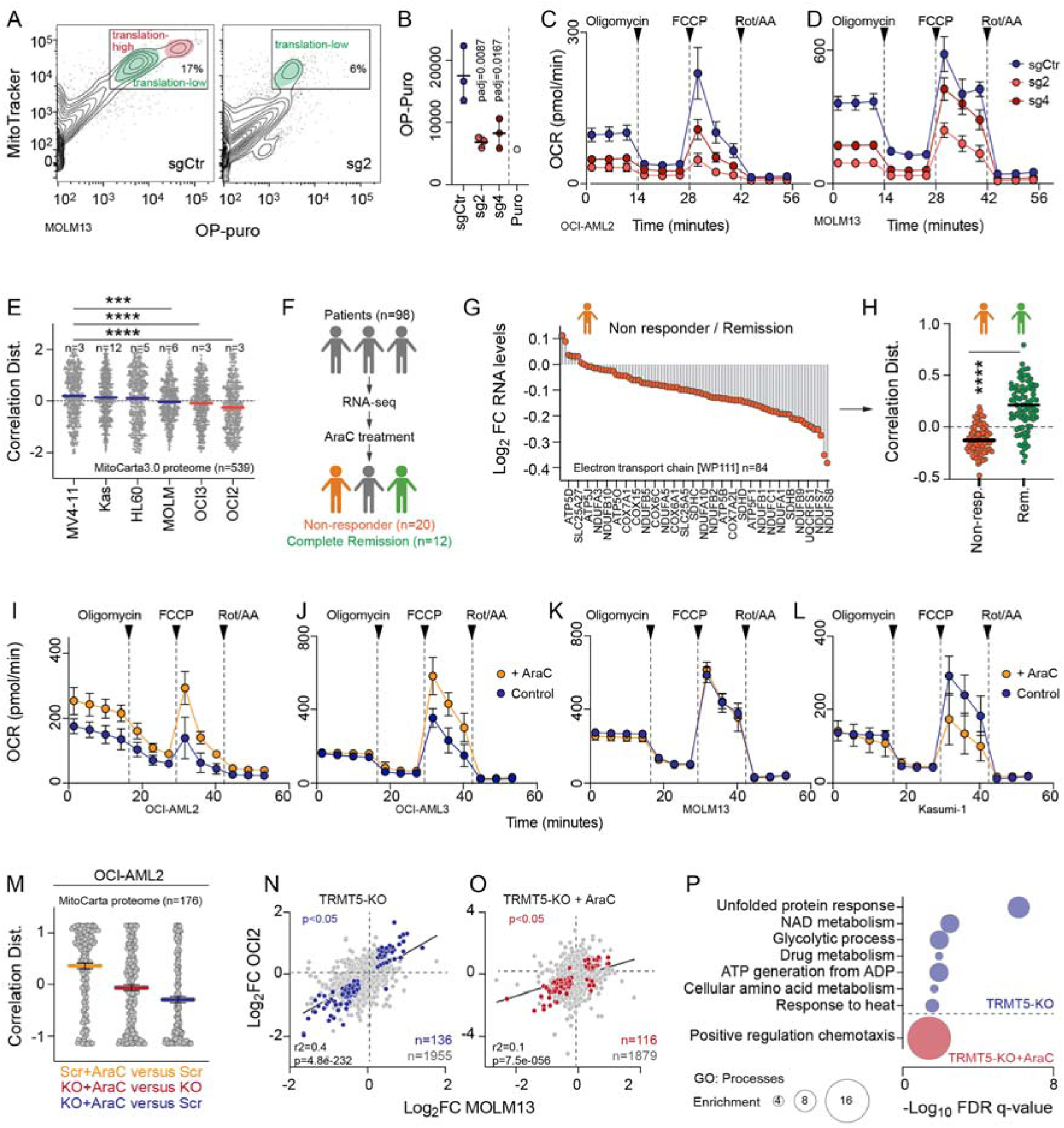
Drug tolerance requires switching from low to high OXPHOS in AML cells. **(A,B)** Flow cytometry (A) and quantification (B) of O-Propargyl-puromycin (OP-puro) incorporation into nascent polypeptides in mitochondria of TRMT5-depleted (sg2, sg4) and control (sgCtr) MOLM13 cells. Puromycin-treated cells served as a control. (**C,D**) Oxygen consumption rate (OCR) in OCI-AML2 (C) and MOLM13 (D) TRMT5-knockout (sg2, sg4) and sgCtr cells. (**E**) Mitochondrial protein levels in the indicated cell lines assessed by mass spectrometry. Shown are proteins present in all cell lines (n=539). Averaged protein abundance was used for clustering using correlation distance as measured (n=number of proteome analyses). (**F-H**) Illustration (F), log_2_ fold change (FC), and correlation distance (H) of RNA levels of gene encoding proteins of the electron transport chain (WP111) in patients responding (green) or not responding (orange) to cytarabine treatment. (**I-L**) OCR in drug-tolerant OCI-AML2 (I), OCI-AML3 (J) and drug-sensitive MOLM13 (K) and Kasumi-1 (L) cells exposed to cytarabine (AraC) treated (orange) or untreated (control) (blue). (**M**) Correlation Distance of log_2_ fold changes of nascent translation of mitochondrial genes (n=176) in AraC-treated control (scr) OCI-AML2 cells (yellow), AraC-treated TRMT5-knockout cells (red) or AraC TRMT5-knockout cells versus control (scr) cells (blue). Scr: Infection of a scramble sgRNA. (**N-P**). Correlation of log_2_ FC of nascent translation of TRMT5 OCI-AML2-(AML2) and MOLM13-knockout cells in the absence (N) or presence (O) of AraC and corresponding gene enrichment analyses (P) (GOrilla). Mean ± SD (B-D,I-L). Mean (E,H,M). Dunnett’s multiple comparisons test (B).

### Drug tolerant cells switch from low to high OXPHOS in acute stress responses

High OXPHOS levels are linked to chemotherapy resistance *in vivo* ^23^, indicating that drug-tolerant AML cells primarily rely on OXPHOS for energy production. Unexpectedly, we found that drug-tolerant cancer cells had on average the lowest levels of mitochondrial protein expression (MitoCarta3.0; n=1136). These cells also showed comparably low levels of basal and maximal oxygen consumption rates (Figure 5E; Supplementary Figure 5C,D; Supplementary Table 8).

To test whether our finding that cultured drug-tolerant leukemic cells displayed low mitochondrial activity was disease-relevant *in vivo*, we analysed RNA-sequencing data from 98 patients obtained at the time of AML diagnosis (Figure 5F) ^25^. After first induction therapy, we analyzed those patients who showed complete remission (n=12), and those who did not respond to AraC treatment (n=20). RNA levels of genes encoding for electron transport chain proteins were consistently lower in non-responding patients (Figure 5G,H; Supplementary Figure 5E,F). We concluded that mitochondrial OXPHOS was not the main energy source fuelling growth of inherently drug tolerant cells.

To directly test how exposure to stress affected mitochondrial activity, we exposed drug-tolerant and -sensitive AML lines to AraC and quantified cellular respiration (Figure 5I-L). Only the drug-tolerant OCI-AML2/3 cells significantly up-regulated oxygen consumption rates (OCR) in response to AraC (Figure 5I,J). In contrast, respiration in AraC-sensitive MOLM13 and Kasumi-1 cells were unaffected or even reduced (Figure 5K,L). Thus, our data indicated that the capacity to up-regulate oxygen consumption rates rather than high OXPHOS steady state levels contributed to drug tolerance. Indeed, exposure to AraC up-regulated mRNA translation of mitochondrial proteins solely in the presence of a functional TRMT5 protein (Figure 5M).

Our data so far showed that cell growth of drug tolerant cells was not primarily fuelled by mitochondria. However, only resistant cells had the capacity to dynamically switch from low to high OXPHOS when exposed to AraC.

### TRMT5 is required for increasing mitochondrial energy efficiency

To determine how TRMT5 regulated cellular survival, we performed nascent proteomics analyses and found that translation of 136 mRNAs significantly changed in both TRMT5 knockout lines (Figure 5N). Proteins regulating the unfolded protein response and energy metabolism pathways were most affected by loss of TRMT5 (Figure 5P, blue circles). Activation of the unfolded protein response can be explained by accumulation of ribosomal +1-frameshifts caused by the elimination of m^1^G37 ^26,27^. An essential role of TRMT5 in regulating mitochondrial respiration has previously been shown in patients carrying loss-of-function *TRMT5* mutations. Hypomethylation of m^1^G37 in mt-tRNAs causes multiple respiratory-chain deficiencies ^21^.

In contrast, alterations in nascent translation only poorly correlated when the two TRMT5-depleted lines were exposed to AraC, confirming that drug-tolerant and -sensitive cells responded very differently to the drug treatment (Figure 5O). The only common change in both lines involved translation of genes regulating chemotaxis (Figure 5P; red circle). We concluded that drug-tolerant and -sensitive cells have distinct energy requirements to fuel cell growth, but up-regulation of mitochondrial metabolic response was strictly required for survival of AraC treatment.

### Drug-tolerant cells withstand inhibition of mitochondria but only in the absence of AraC

To directly test whether drug-tolerant cells required mitochondria for proliferation, we performed growth competition assays in the absence of TRMT5 (Figure 6A). While the number of TRMT5-knockout cells was reduced in both lines (Figure 6B-C), the apoptosis rate was about four-fold higher in drug-sensitive MOLM13 cells (Figure 6D; Supplementary Figure 6A). To track the fate of TRMT5-knockout cells over time, we quantified the number of cells containing the CRISPR-induced mutations over a period of 20 days (Figure 6E, F). OCI-AML2 tolerated TRMT5 mutations for up to 17 days. In contrast, MOLM13-knockout cells were quickly eliminated from the culture (Figure 6E, F).

**Figure 6.**
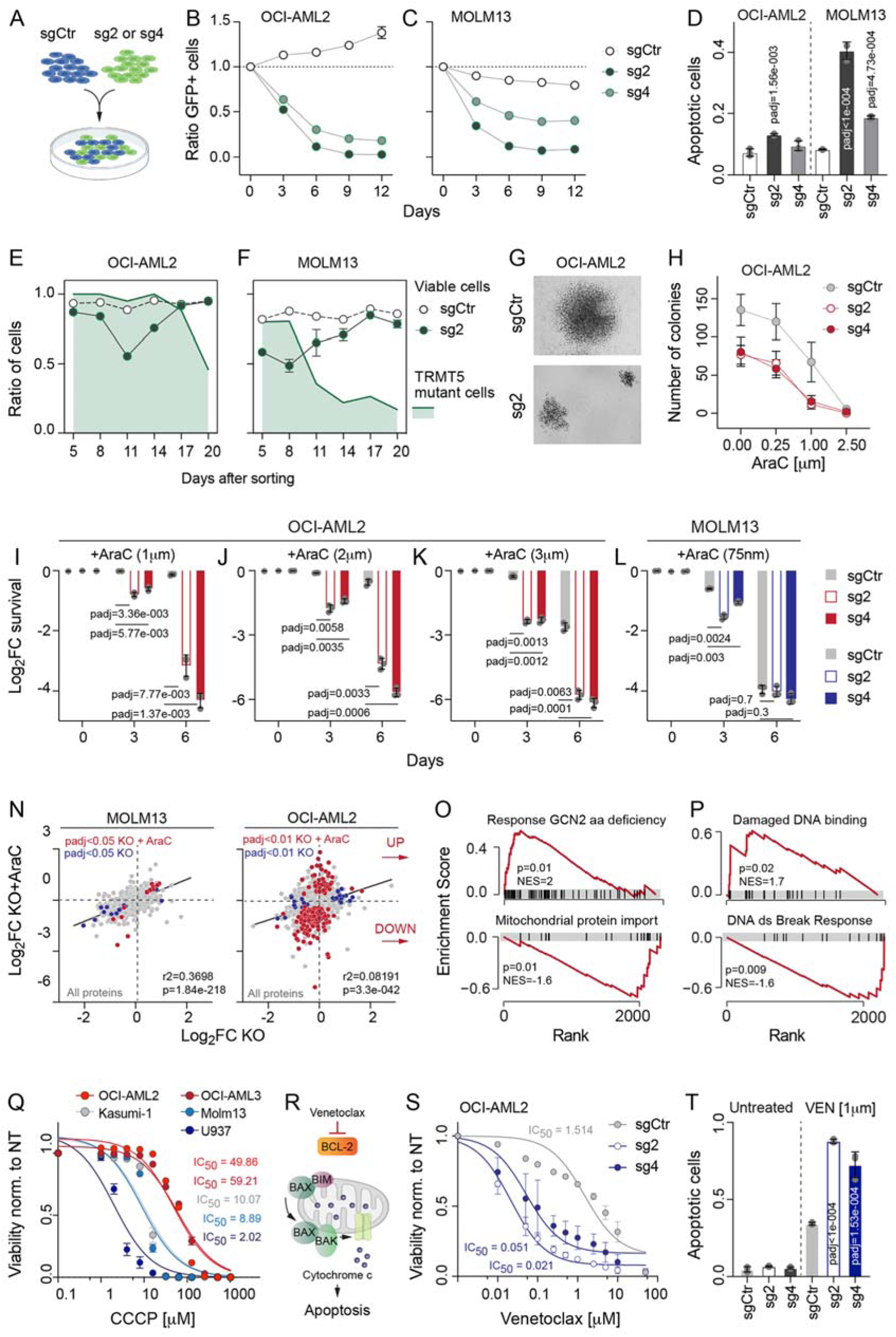
Dynamic mitochondrial activity determines drug response. (**A-C**) Illustration (A) of growth competition assays using GFP-labelled OCI-AML2 (B) and MOLM13 (C) control (sgCtr) and knockout cells (sg2, sg4). (**D**) Frequency of apoptotic cells in populations of OCI-AML2 and MOLM13 control (sgCtr) and knockout (sg2,sg4) cells using Annexin V labelling. (**E,F**) Quantification of mutated OCI-AML2 (E) and MOLM13 (F) cells (green line) overlayed with the frequency of viable TRMT5-knockout (sg2) (black dots) and control (sgCtr) (white dots) cells over time. (**G,H**) Representative bright field image (G) and colony-forming efficiency (CFE) assays (H) of OCI-AML2 control (sgCtr) and knockout (sg2, sg4) cells in the presence of increasing concentrations of AraC. (**I-L**) Log_2_ fold change (FC) of survival of drug-tolerant OCI-AML2 (I-K) and drug-sensitive MOLM13 (L) control (sgCtr) and TRMT5-knockout (sg2, sg4) cells at the indicated time points. (**N**) Correlation of log_2_ FC of nascent translation of MOLM13 (left) and OCI-AML2 (right) TRMT5-knockout (KO) cells compared to cells infected with a scramble guide RNA in the presence or absence of AraC. Blue dots: significantly changed in absence of AraC. Red dots: Significantly changed in the presence of AraC. (**O,P**) Gene set enrichment analysis of significantly up-regulated (top) and down-regulated (bottom) mRNA translation of AraC treated OCI-AML2 KO cells as shown in (N,right panel). (**Q**) Dose-response curve examining the effect of different concentrations of Carbonyl cyanide m-chlorophenyl hydrazone, Carbonyl cyanide 3-chlorophenylhydrazone (CCCP) in the indicated blood cancer lines with corresponding IC50 values. (**R-T**) Illustration of venetoclax mechanism of action (R), IC50 curves (S) and quantification of apoptotic cells of OCI-AML2 control (sgCtr) and KO (sg2,sg4) cells in the presence of venetoclax. Mean ± SD (B-D,H-L,Q,S,T). Dunnett’s multiple comparisons test (D,T). Two-way ANOVA multiple comparisons test (I-L).

To test viability on the single cell level, we performed colony forming unit (CFU) assays. Drug-sensitive MOLM13 cells were unable to form colonies in the absence of TRMT5 (Supplementary Figure 6C). OCI-AML2 knockout cells formed colonies, yet their size and number were consistently reduced both in the absence and presence of AraC (Figure 6G, H). However, as shown in the synthetic lethality screens, treatment with AraC further significantly reduced survival of drug-tolerant TRMT5-knockout cells (Figure 6I-K; Supplementary Figure 6C-E). Drug-sensitive MOLM13 cells on the other hand exhibited equally high levels of apoptosis when exposed to AraC or when TRMT5 was deleted (Figure 6L; Supplementary Figure 6F).

In line with our finding that AraC-treatment and TRMT5-deletion both increased cell death to a similar extent, we did not observe any additional nascent proteomic alterations in MOLM13 knockout cells exposed to AraC (Figure 6N; left panel). In contrast, drug-tolerant OCI-AML2 knockout cells exhibited major changes in mRNA translation in response to AraC-treatment (Figure 6N; right panel). Expression of proteins regulating mitochondrial import were significantly reduced, while levels of stress response proteins were increased (Figure 6O).

In summary, drug-tolerant cells resisted loss of TRMT5 due to low mitochondrial activity in normal growth conditions. However, TRMT5-driven mitochondrial mRNA translation was strictly required for cellular survival in the presence of AraC.

### Mitochondrial energy efficiency determines cellular responses to cytotoxic drugs

Cytarabine is incorporated into DNA and causes DNA damage by stalling replication forks. Accordingly, we measured significant alterations in mRNA translation of genes encoding proteins involved in the DNA damage response (Figure 6P). Since mitochondria sense fluctuations in DNA damage signalling pathways, we asked whether mitochondria only indirectly impacted cell survival by responding to DNA damage in the nucleus when exposed to AraC.

Two lines of evidence suggested that dynamic alteration of mitochondrial activity directly regulated cellular survival. First, both drug-tolerant OCI-AML2/3 cancer lines survived an over 10-fold higher concentration of CCCP, a chemical inhibitor of oxidative phosphorylation, confirming low dependency on OXPHOS under normal growth conditions (Figure 6Q). Second, cell death in response to the BCL2-specific inhibitor venetoclax also increased in TRMT5-depleted cells (Figure 6S,T). In conclusion, TRMT5-dependent mitochondrial flexibility directly mediated cellular survival in response to chemotherapeutic drugs.

In summary, we propose that mitochondrial inhibitors can circumvent adaptive resistance to venetoclax and cytarabine therapy in acute myeloid leukemia.

## Discussion

Here, we performed targeted functional genetic screenings to identify novel RNA modification pathways involved in cancer cell growth and drug resistance. The currently best-studied modification in cancer development and progression is N6-methyladenosine (m^6^A), the most abundant internal mRNA modification. We now substantially expand the list of cancer-relevant RNA modifications by showing that at least 50 out of the 150 known RNA modifying proteins were essential in at least one of the 18 tested cell lines. RNA modifying proteins universally required for cell survival equally targeted mRNA, tRNA and rRNA species, highlighting the importance of both transcriptional and translational processes to maintain basic cellular functions.

Expression levels of RNA modifying proteins were on average higher in tumour cells when compared to their healthy counterpart, yet we find no evidence for essential tissue-specific RNA modifications. Low tissue-specificity can be explained by the high number of rRNA- and tRNA-modifying enzymes in our tested gene pool which are expected to be expressed in most cells. The most extensive modified type of RNA is cytosolic and mitochondrial tRNA containing more than 20 distinct modifications mediated by about 40 regulatory proteins. Together, tRNA modifications enhance mRNA translation speed accuracy and fidelity and in addition, control mitochondrial energy efficiency to fuel protein synthesis. We now show that disturbing this regulatory network by depleting single RNA modifications can have far reaching consequences for cellular survival and drug resistance.

Unexpectedly, the number of essential RNA modifying proteins varied substantially between the cell lines even within the same cancer type such as leukemia. We provide evidence that the high level of variability of essential genes between cell types reflected inherent metabolic requirement needed to fuel optimal cell growth and survival in response to stress signals. Leukemic cell lines with comparably low levels of OXPHOS tolerated the depletion of mitochondrial RNA modifying enzymes better than cells relying on mitochondrial energy production to fuel cell growth. Surprisingly, those cells were also inherently drug tolerant. However, synthetic lethal screens identified a novel vulnerability of drug tolerant AML cells. To survive drug treatments, cells with low OXPHOS must up-regulate mitochondrial energy production. Interrupting mitochondrial mRNA translation by depleting mitochondrial RNA modifications such as TRMT5-mediated m^1^G formation in the tRNA anticodon loop re-sensitized the cells towards chemotherapeutic drug treatments.

In summary, by performing CRISPR-based genetic screening targeting 150 RNA modifying proteins in cancer cells, we identified novel cancer-relevant essential RNA modifications, and we reveal TRMT5 as promising novel drug target in the treatment of chemotherapy-resistant acute myeloid leukemia.

## Material and methods

### Cell culture

Primary cell lines normal human epidermal keratinocytes (NHEK) and normal human epidermal melanocytes (NHEM) were purchased from PromoCell. The oral squamous cell carcinoma cell line VDH15 was generated as previously reported ^28^. All other cancer cell lines were ordered from DSMZ or ATCC.

The leukaemia cells lines U937, HL60, Kasumi-1, MOLM13 were cultured in RPMI-1640 (Sigma-Aldrich) supplemented with 10% (U937, HL60) or 20% (Kasumi-1, MOLM13) fetal bovine serum (FBS, Bio&SELL GmbH, Feucht, Germany) and 1% penicillin-streptomycin (PS, Thermo Fisher Scientific, 10.000 U/ml). For the cell lines OCI-AML2 and OCI-AML3 the medium MEM alpha (Thermo Fisher Scientific) supplemented with 20% FBS, and 1% Penicillin-Streptomycin (PS) was used.

The lung cancer cell lines A549 and PC-9 were kept in RPMI-1640 with 10% FBS and 1% PS. Primary melanocytes (NHEM) were kept in melanocyte growth medium M3 (PromoCell) with 1% PS. A2058 and A357 were grown in DMEM (Thermo Fisher Scientific) with 10% FBS and 1% PS. The HNSCC cell lines Cal33 and Fadu were grown in DMEM with 10% FBS and 1% PS while SCC25 was cultured in DMEM/F12 (Thermo Fisher Scientific) with 15% FBS and 1% PS. VDH15 was cultured in defined keratinocyte SFM (Thermo Fisher Scientific) with 1% PS. The NHEK cells were grown in keratinocyte growth medium 3 (PromoCell) with 1% PS. OPM2 and RPMI-8226 cells are derived from Myeloma cells and were kept in RPMI-1640 with 10% FBS, 1% GlutaMax (Thermo Fisher Scientific, Germany) and 1% PS. Cells were split every 2 to 3 days and cell concentrations were kept between 0.5 and 2 Mio cells per ml. The HEK293 T-derived cell line Lenti-X (Takara) was cultured in DMEM with 10% FBS and 1% PS. Suspension and adherent cells were cultured in a humidified incubator at 37°C with 5% CO2. Prior starting with the CRISPR-Cas9 based screens cell lines were authenticated by STR-profiling and cells were tested for mycoplasma contamination.

### Targeted CRISPR Cas9 genetic screening

An RNA-modification-focused CRISPR-Cas9 library consisting of 1200 single-guide RNAs (sgRNA) against 150 RNA-modifying proteins and around 400 sgRNAs against the 250 most abundant snoRNAs in AML was designed. Further, sgRNAs against 8 control genes (controls) as well as 50 nontargeting sgRNAs were included. The total 1708 library sgRNAs were cloned into the lentiCRISPRv2-GFP vector (82461, Addgene). The sgRNA-bearing plasmids were transduced together with the lentiviral envelope plasmid pDM2g (12259, Addgene) and the packaging plasmid psPAX2 (12260, Addgene) into lenti-X 293T cells using TurboFect transfection reagent (R0531, Thermo Fisher Scientific). Lentivirus was harvested after 72 hours and concentrated via ultracentrifugation. Before infection, virus concentration was titrated to aim for a multiplicity of infection (MOI) of 0.3 to guarantee that a single cell is infected by one virus.

The cancer cell lines and the healthy human control cell lines were infected and the input samples were taken after 48 hours which marked day 1 of the experiment. Samples were picked on days 8 and 22. In addition to the broad screening of the different cancer cell lines, the leukemia cell line OCI-AML2 was sorted for GFP 96 hours after infection and then divided into a treatment and a control group. The treatment group was stimulated for 3 days with cytarabine (STADA) and then 4 days were given for recovery. Three treatment-recovery cycles were performed with increasing doses of cytarabine (1µM, 2µM, 3µM). Again, samples were taken on days 8, 15, and 22.

The whole genomic DNA of the samples was isolated with the Quick-DNA Midiprep Plus Kit (Zymo Research). To ensure a coverage of 500x we aimed for a total PCR-input amount of 20µg DNA in the unsorted and of 10µg DNA in the sorted cell populations. The preparation and sequencing of the sgRNAs was similarly performed as described before ^29,30^. The sequencing primers were ordered in NGS-grade quality from Eurofins. The primers sequences are as follows: 1^st^ primer forward (AAT GGA CTA TCA TAT GCT TAC CGT AAC TTG AAA GTA TTT CG), 1^st^ primer reverse (TCT ACT ATT CTT TCC CCT GCA CTG TTG TGG GCG ATG TGC GCT CTG), 2^nd^ primer forward (AAT GAT ACG GCG ACC ACC GAG ATC TAC ACT CTT TCC CTA CAC GAC GCT CTT CCG ATC T—1-9bp variable sequence—8bp barcode—TCT TGT GGA AAG GAC GAA ACA CCG), 2^nd^ primer reverse (CAA GCA GAA GAC GGC ATA CGA GAT—8bp barcode—GTG ACT GGA GTT CAG ACG TGT GCT CTT CCG ATC T—1-9bp variable sequence—TCT ACT ATT CTT TCC CCT GCA CTG T).

In a first nested Polymerase Chain Reaction (PCR) approach, the sgRNA encoding sequences were amplified from the vector backbone. A subsequent PCR added the Illumina overhangs and a specific pair of barcoding sequences to the amplified sgRNAs for multiplexing. PCR products were purified with AMPure XP beads (A63881, Beckmann Coulter) and the quality was checked using an Agilent Bioanalyzer System. The DNA libraries were sequenced using an Illumina NextSeq500 instrument for next-generation sequencing. Reads were demultiplexed using the P7 barcode in addition to the in-line barcode information. Each read was scanned for the vector motif and the subsequent 20 nucleotides were used to assign one of the 1708 unique sgRNA from the screen. The robust ranking aggregation (RRA) algorithm in MAGeCK software v0.5.9.4 was used to identify positively or negatively enriched genes.

### Library generation and RNA sequencing

For RNA sequencing experiments, RNA was extracted from cells using Direct-zol RNA Miniprep Kits (Zymo Research) and libraries were generated using the Illumina Total RNA Prep with Ribo-Zero Plus kit following manufacturers’ instructions. Samples from the different cell lines were multiplexed and sequenced on a NovaSeq 6000 PE 100bp S4 sequencing platform (Illumina). Three replicates per cell lines were sequenced.

### RNA sequencing analyses

RNA sequencing data were processed by the DKFZ One Touch Pipeline (OTP) ^31^ using the RNA-seq workflow version 1.3.0 (https://github.com/DKFZ-ODCF/RNAseqWorkflow) in combination with the workflow management system Roddy version 3.5.8 or 3.5.9 (https://github.com/TheRoddyWMS/Roddy). In brief, data were aligned against the appropriate reference genome: 1KGRef_PhiX (generated from the 1000 Genomes assembly, based on hs37d5 and including decoy sequences merged with PhiX contigs to be able to align spike in reads) using the STAR aligner version 2.5.3a ^32^. Duplicate marking was performed using Sambamba version 0.6.5 ^33^ and quality control was performed using Samtools version 1.6 flagstat ^34^ as well as rnaseqc version 1.1.8 ^35^. FeatureCounts of the Subread package version 1.5.1 ^36^ was used for gene specific quantification of reads on the appropriate gene annotation: GENCODE version 19 or GENCODE version M12 in the strand-specific counting mode.

Raw gene count values were then used as input for a differential gene expression analysis using DESeq2 version 1.28.1 ^37^ within R version 4.0.0 R Core Team (2020). R: A language and environment for statistical computing. Foundation for Statistical Computing, Vienna, Austria. https://www.R-project.org/), which was made using default conditions besides fitType=“local” for each experiment separately. Results tables were generated for each treatment compared to the appropriate control.

### Generation of single CRISPR Cas9-knockout cells

For a single knockout of TRMT5 two sgRNAs (sgRNA2, sgRNA4) of the library pool were selected. Infected cells were sorted after 72 to 96h for GFP. Successful knockout was validated via Western blot. Experiments were performed 5 to 14 days after sorting.

### Western blotting

Cells (2 – 5 x 10^6^) were washed two times with cold PBS and lysed with RIPA buffer at 4°C (Thermo Fisher Scientific) supplemented with protease inhibitor (Roche) and optional phosphatase inhibitor (Roche.). Cell lysates were centrifuged and the supernatant was stored at −20 degrees. Protein concentration was measured via the Pierce BCA Protein Assay kit (Thermo Fisher Scientific) according to the manufacturer’s instructions and measured using a spectrophotometer (Tecan, Spark®). Between 20 to 50 ng of protein (for each run the same amount per sample) was added to 4X LDS sample buffer (Thermo Fisher Scientific) and 10X sample-reducing agent (Thermo Fisher Scientific), boiled for 3 minutes at 95°C and loaded on a 4% to 12% (alternatively 4% to 20%) tris-glycine gradient gel (Thermo Fisher Scientific) for sodium dodecyl sulfate polyacrylamide gel electrophoresis (SDS-PAGE). Proteins were blotted onto a nitrocellulose membrane for 90 minutes with a current intensity of 350 mA. The nitrocellulose membrane was blocked for 45 minutes in 5% (w/v) non-fat milk or bovine serum albumin diluted in 1x TBS, and 0.1% Tween-20 (TBS-T) and incubated with the primary antibody in a blocking solution at 4°C overnight.

After washing each membrane 3 times for 5 min with TBS-T, anti-rabbit or mouse horseradish peroxidase (HRP)-labeled secondary antibodies (1:4000, P044701-2 and P044801-2, Agilent) were added for 1 hour at RT. Membranes were washed 3 times again and the chemiluminescence signal was captured by using ECL reagents (RPN2232, GE Healthcare; 34095, Thermo Fisher Scientific). The following primary antibodies were used: anti-TRMT5 (1:1000, ab259986, Abcam) and anti-GAPDH (1:2000, 2118, Cell Signaling).

### Cell competition assay

Equal numbers of GFP-expressing TRMT5-depleted or scramble infected cells (1 x 10^5^) were seeded together and grown in 750 µl of medium (24 well plate). Samples were taken every 3 days for flow cytometry analysis.

### Colony-forming unit (CFU) assay

80 ml of cytokine-free methylcellulose for human cells (H4230, StemCell^TM^) was mixed with 20 ml of medium and 1 ml of penicillin and streptomycin. Optionally, a determined amount of cytarabine was added to the methylcellulose mix. 300 cells were added to the 600µl methylcellulose and were seeded onto a 12-well plate. Colonies were counted after 10 days of incubation.

### Annexin apoptosis assay

Infected cells (3 x 10^5^) were seeded in 1 ml of an appropriate medium onto a 12-well plate. Depending on the predetermined IC50 of the wild-type cell line, cells were stimulated with 2 to 3 increasing doses of cytarabine. After 48 hours, 200µl of cell suspension was taken and stained with 0.75 µl Annexin V pacific blue (A35122, Thermo Fisher Scientific) and 0,25 µl Propidium Iodide (BD Biosciences) in 100 µl Annexin V buffer (V13246, Thermo Fisher Scientific). Apoptotic and necrotic cells were measured via flow cytometry. The remaining cells were seeded and stimulated again with the same drug dose. After 72 hours, the assessment of apoptotic and necrotic cells was repeated.

### Sanger sequencing and TIDE sequencing

Infected and GFP-sorted cells were cultured for 25 days. Starting at day 5, cell viability was measured every 3 days using annexin V/PI staining (see above). In addition, samples of 5 x 10^4^ cells were taken at the same time points. Genomic DNA was isolated using the QIAamp UCP DNA Micro Kit (Qiagen). PCR was performed to amplify the TRMT5 gene (forward primer: GTC AGC ATT TAG CCC GTT ATT TGA G, reverse primer: CTG AAG GCC TAA CAA GTC TTG TGT AC). The PCR product was purified via gel electrophoresis and recovered using the Zymoclean Gel DNA Recovery Kit (ZymoResearch). 50 ng of the purified DNA was sent to Eurofins Genomics for Sanger sequencing (sequencing primer: GCC ATT TGG ATT CTC AGG AAG). Sequencing results were analyzed via TIDE an open-source software to track Indels by computational decomposition ^38^.

### OP-Puromycin incorporation into mitochondrial nascent polypeptides

Cells were seeded at a density of 6 x 10^5^ cells per ml one day before the experiment. O-propargyl-puromycin (Thermo Fisher Scientific) with a final concentration of 50 µM and MitoTracker Deep Red (Thermo Fisher Scientific) with a final concentration of 200 nM were added for an incubation of 45 minutes. For positive control, one sample was treated with 1 mM of puromycin. Mitochondria were isolated using the Mitochondria Isolation Kit for mammalian cells (Thermo Fisher Scientific). Isolated mitochondria were fixed with 4% paraformaldehyde for 15 minutes and permeabilized according to the Click-iT™ Cell Reaction Buffer Kit (Thermo Fisher). O-propargyl-puromycin detection was performed according to the manufacturer’s instruction. Samples were analyzed via flow cytometry (BD-Bioscience Fortessa).

### Seahorse assay

Infected cells were seeded at a density of 6 x 10^5^ cells per ml one day before the experiment. Oxygen consumption rates (OCR) were measured on a Seahorse XFe96 extracellular flux analyzer (Agilent) following the manufacturer’s instructions. Briefly, AML suspension cells were collected from culture plates and seeded into 96 wells PDL coated plate (Agilent) at a concentration of 1.5 x 10^5^ cells per well, followed by centrifugation to attach the cells to the bottom of the well. Plates were equilibrated for 1 hour in XF assay medium supplemented with 10 mM glucose, 1 mM sodium pyruvate, and 2 mM glutamine in a non-CO2 incubator. OCR was monitored at baseline and throughout sequential injections of oligomycin (1 μM), carbonyl cyanide-4-(trifluoromethoxy) phenylhydrazone (1 μM) and rotenone/antimycin A (0.5 μM each). Shaking events were reduced to 30 seconds to avoid detachment of cells from the plate. Data for each well were normalized to protein concentration as determined using Pierce BCA Protein Assay kit (ThermoFisher) after the measurement on the XFe96 machine.

### Total proteomics

For total proteome analyses of different cell lines, cells were grown at equal density overnight. GFP-sorted TRMT5-knockout and control (scramble) cells were harvested on days 12 and 13 after infection. 4×10^6^ cells per condition and replicate were washed thrice with ice-cold 1x PBS. Experiments were conducted at least in triplicates.

Cells were lysed in 50 mM ammonium bicarbonate containing 0.1% Rapigest (Waters) and sonicated samples were reduced with dithiothreitol, alkylated with chloroacetamide and digested with trypsin and rLysC (Promega). After acidification samples were spun and supernatants were used for injections. A peptide amount corresponding to 1 μg was analyzed on a TriLJHybrid Orbitrap Fusion mass spectrometer (Thermo Fisher Scientific) operated in dataLJdependent acquisition mode with HCD fragmentation. RAW data were processed with MaxQuant (1.6.2.6) ^39^ using default settings. Only proteins with at least 1 unique peptide were considered as identified and normalized LFQ values were used for quantitative comparative analyses. The gene set enrichment analyses (GSEA) were performed using R package fgsea ^40^ (version 1.6.0) with a *P*-value ranking of proteins, gene sets defined by the REACTOME pathway database (R package ReactomePA version 1.24.0) ^41^. Proteome data was further analyzed using the MitoCarta3.0 inventory ^42^.

### Nascent proteomics

Nascent proteomics studies were performed as described before ^43^. In case of drug treatment, MOLM13 cells were treated with 15 nM AraC and OCI-AML2 cells were treated with 1μM AraC for 16 hours. To deplete cells of methionine, lysine, and arginine, the cells were then incubated for 30 min in depletion medium (medium without methionine, arginine, and lysine). Cells were pulse-labeled for 6 hours with AHA- and SILAC-amino acid containing medium. Compared conditions were labeled with different SILAC medium (heavy (HV) or intermediate (IM) l-arginine and l-lysine, Cambridge Isotope Laboratories, Inc., Tewksbury). In case of previous drug treatment, the respective drug concentration was also added to the labeling medium. After harvest, washing and cell lysis, equal amounts (750 μg each) from different conditions and SILAC labels ((HV)/(IM)) were combined and proceeded to AHA-enrichment. To exclude label-specific effects replicates were grown in switched HV/IM conditions. Experiments were conducted at least in triplicates. Newly synthesized proteins were enriched using the Click-iT^®^ Protein Enrichment Kit (Invitrogen, Schwerte, Germany), applying the vendor’s protocol with slight modifications as described. A peptide amount corresponding to 1 μg was analyzed on a TriLJHybrid Orbitrap Fusion mass spectrometer (Thermo Fisher Scientific) operated in dataLJdependent acquisition mode with HCD fragmentation. RAW data were processed with MaxQuant.

### Small mitochondrial RNA isolation

Mitochondria were isolated using a MACS antibody based mitochondria extraction kit (Miltenyi Biotech) to obtain high quality and intact mitochondria for subsequent RNA isolation. Subsequently, small RNAs were isolated using the *mir*Vana™(Thermo Fisher Scientific) or miRNeasy Micro Kit (Qiagen).

### Mass Spectrometry for quantifying RNA modifications levels

tRNA was enzymatically digested using benzonase (Santa Cruz Biotech) and nuclease P1 (Sigma) in 10 mM ammonium acetate pH 6.0 and 1 mM MgCl2 at 40 °C for 1 h, added ammonium bicarbonate to 50 mM, phosphodiesterase I and alkaline phosphatase (Sigma) and incubated further at 37 °C for 1 h. Digested samples were precipitated with 3 volumes of acetonitrile and supernatants were lyophilized and dissolved for LC-MS/MS analysis. The mobile phase consisted of A: water and B: methanol (both added 0.1% formic acid) run at 0.22 ml/min, starting with 5% B for 0.5 min followed by 2.5 min of 5-20% B, 3.5 min of 20-95% B, and 4 min re-equilibration with 5% B. Mass spectrometric detection was performed using an Agilent 6495 Triple Quadrupole system monitoring the mass transitions 268.1-136.1 (A), 284.1-152.1 (G), 244.1-112.1 (C), 245.1-113.1 (U), 298.1-166.1 (m^1^G), 413.1-281.1 (t^6^A), 301.1/152.1 (d_3_-Gm), 285.1/153.1 (d_3_-m^6^A), 273.1/136.1 (^13^C_5_-A), 246.1/114.1 (d_2_-C) in positive ionization mode.

### Nanopore tRNA library preparation, sequencing and data analysis

To sequence native tRNAs using nanopore sequencing, we employed Nano-tRNAseq. Briefly, small RNA fractions isolated from TRMT5-knockout and control samples were deacylated in 100 mM Tris-HCl (pH 9.0) at 37°C for 30 min. Deacylated tRNAs were recovered using Zymo RNA Clean and Concentrator kit (Zymo, R1016), were then ligated to pre-annealed Nano-tRNAseq 5’ and 3’ RNA adapters at room temperature for 2 hours with 20% PEG PEG 8000 (NEB, B10048) using recombinant *E. coli* T4 RNA 2 ligase (made in-house), and followed by clean up with RNAClean XP beads (Beckman, A63987). RNA concentration was determined using Qubit™ RNA High Sensitivity (HS) assay kit (Thermo, Q32852). 50 ng of clean RNA product was then ligated to the RTA adapter (ONT, SQK-RNA002) using concentrated T4 DNA Ligase (NEB, M0202M, 2,000,000 units/mL) at room temperature for 10 minutes, reverse transcribed using Maxima H Minus Reverse Transcriptase (Life Technologies, EP0751), and cleaned up with RNAClean XP beads following manufacturer’s instructions. Then, 6 μl ONT RMX sequencing adapters (ONT, SQK-RNA002) were ligated at room temperature for 10 minutes in a total reaction volume of 40 µl with 1X Quick Ligation Reaction buffer, 3 μl T4 DNA Ligase. The ligation mix was cleaned up with 2X volume of RNAClean XP beads, and washed twice with 150 μl WSB (Wash Buffer, SQK-RNA002). The sample was eluted in 20 μl ELB (Elution Buffer, SQK-RNA002) and incubated for 10 minutes at room temperature on a Hula mixer. The final library was prepared by adding 17.5 μl of nuclease-free water and 37.5 μl of vortexed RRB, and was loaded onto a MinION R9.4 flowcell (FLO-MIN-106).

Reads were base-called using Guppy v3.6.1 in high-accuracy (hac) mode. All Us were converted to Ts, Reads were mapped using BWA (arXiv:1303.3997v2) with parameters *bwa mem -W13 -k6 -xont2d -T20* to the mature human tRNA reference set, obtained from gtRNAdb 2.0 ^44^, to which we added the ligated adapter sequences. Differential tRNA modifications were analysed using by extracting basecalling “errors” such as mismatch, insertion and deletion for each position, using the *get_sum_err.py* script. The output of this script was then used to calculated differential “errors” between WT and TRMT5 KO samples, which were visualized using the *plot_heatmap.py* script. Both scripts are publicly available in Github (https://github.com/novoalab/Nano-tRNAseq).

## Data availability

The CRISPR/Cas9 genetic screening data are available as SinyApp on: https://shiny-portal.embl.de/shinyapps/app/15_rnamod_trmt5. RNA sequencing data using VDH15 cells are available on EGA under the accession number EGAS00001004765 including the attached study EGAD00001010172. All other sequencing data are deposited on GEO under the accession number GSE228208. Nanopore tRNA sequencing data (base-called FASTQ and counts) have been deposited in Gene Expression Omnibus (GEO) under accession code GSE227817. Quantitative proteomics raw data are available in Supplementary Tables 7 to 9a-h. Results are in part based on Cancer Cell Line Encyclopedia ^17^ and pan-cancer analysis of whole genomes (ICGC/TCGA) ^16^ and were downloaded from https://www.cbioportal.org/.

---

**Supplementary Figure 1.**
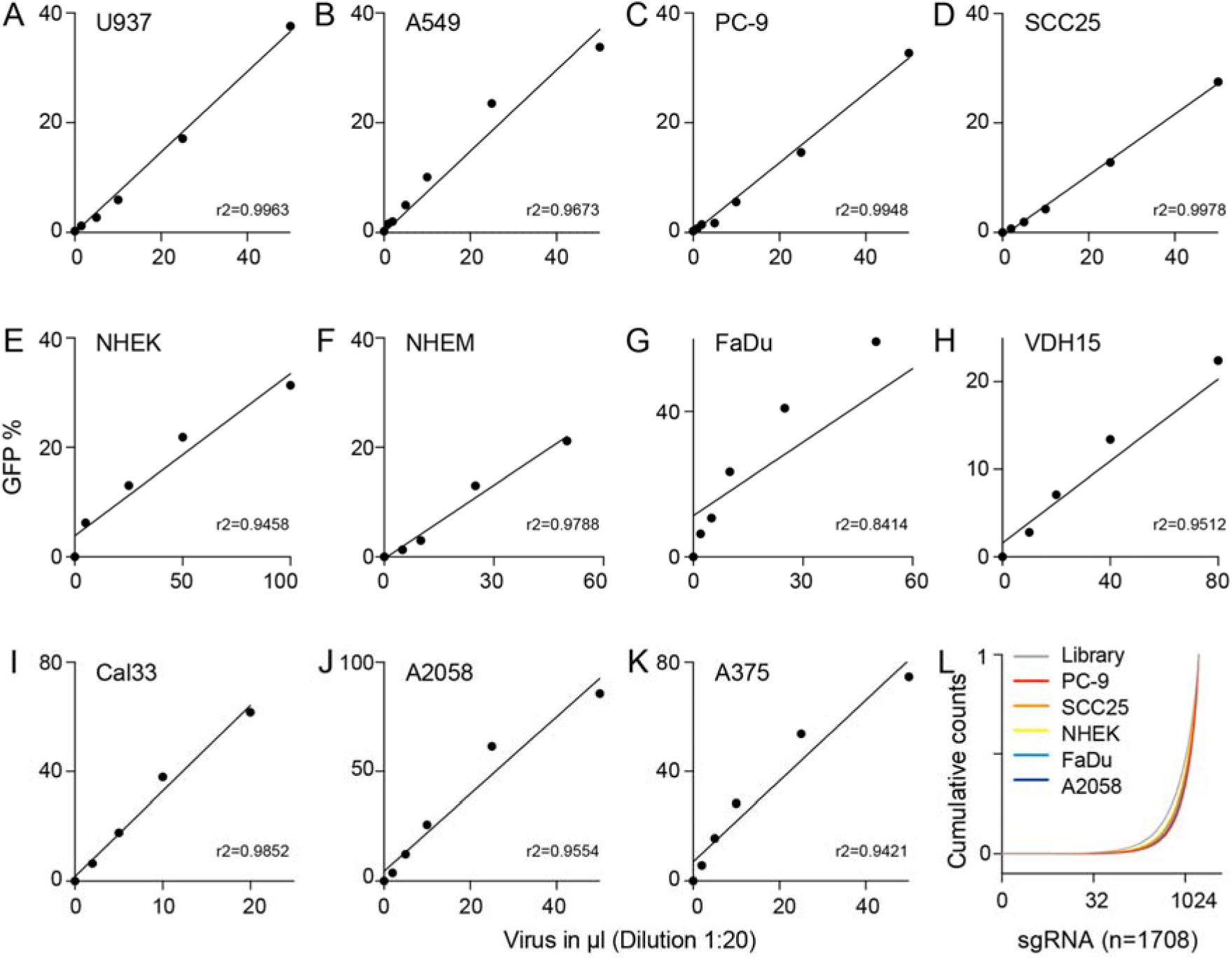
Quality checks of targeted CRISPR/Cas9 screen. (**A-K**) Virus titration in to a MOI of 0.3 in the indicated cell lines. (**L**) Cumulative sgRNA sequencing read counts in the indicated cell lines and in the library (grey).

**Supplementary Figure 2.**
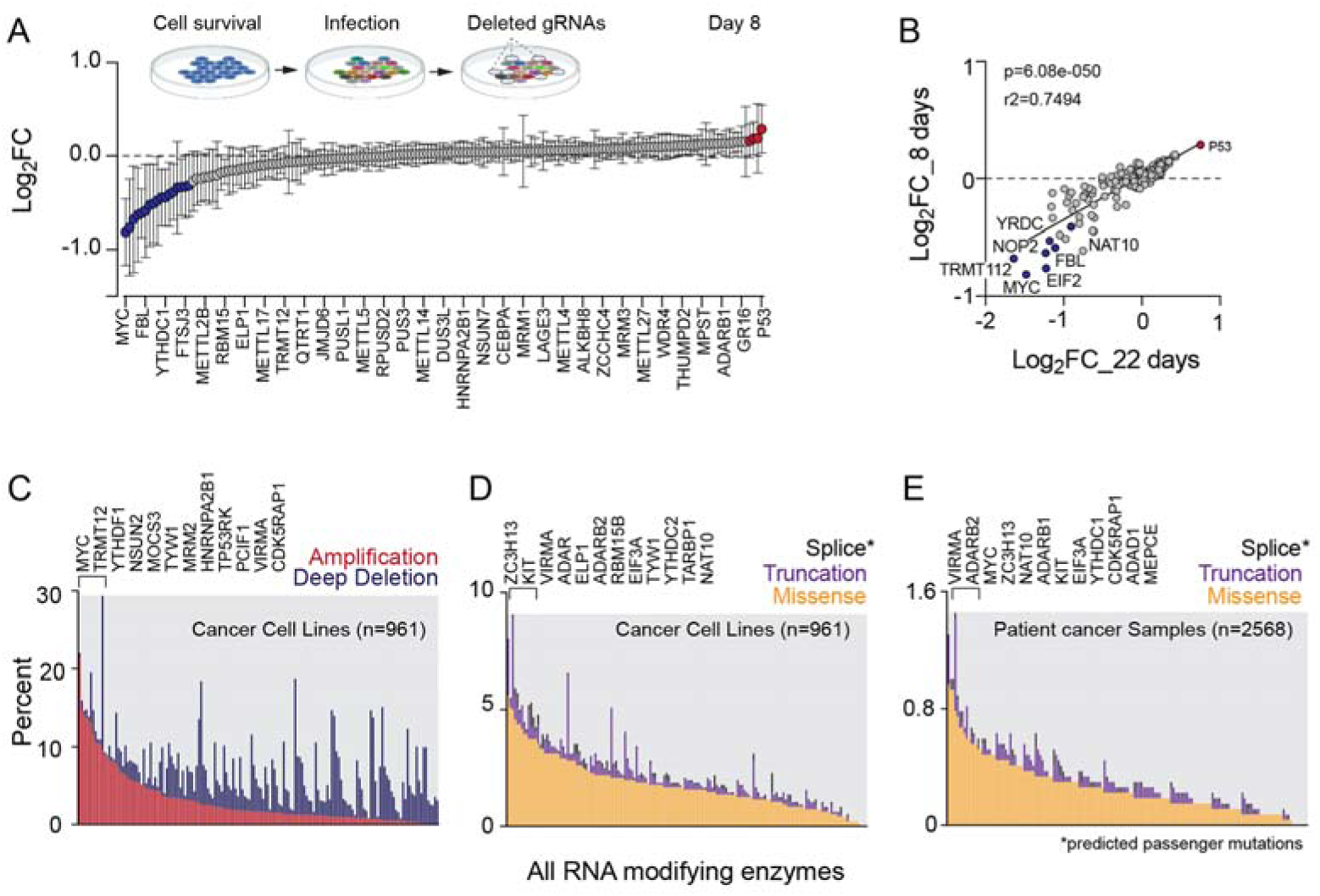
Essential and pan cancer-relevant RNA modifying proteins. **(A)** Scheme of dropout screen (top) and average log_2_ fold-change (FC) of 150 human RNA modifying proteins (RMPs) including positive controls in 18 cell lines (bottom). Shown is every sixth RMP eight days after library infection. (**B**) Correlation between log2 fold-change (FC) of gRNA targeting 150 RMPs including controls at day eight and 22 of the experiment. (**C-E**) Percentage of cancer cell lines (C,D) or tumour samples (E) showing genetic amplification (red) or deep deletions (blue) (C) or truncation (purple) or missense (yellow) mutations (D,E) in 150 RMPs genes including positive controls. RMPs were covered in 961 cancer cell lines (C,D) and 2568 cancer samples spanning 38 different tumour types (E) (cBioPortal). Top 12 amplified RMPs are indicated (top). Mean ± SD (A).

**Supplementary Figure 3.**
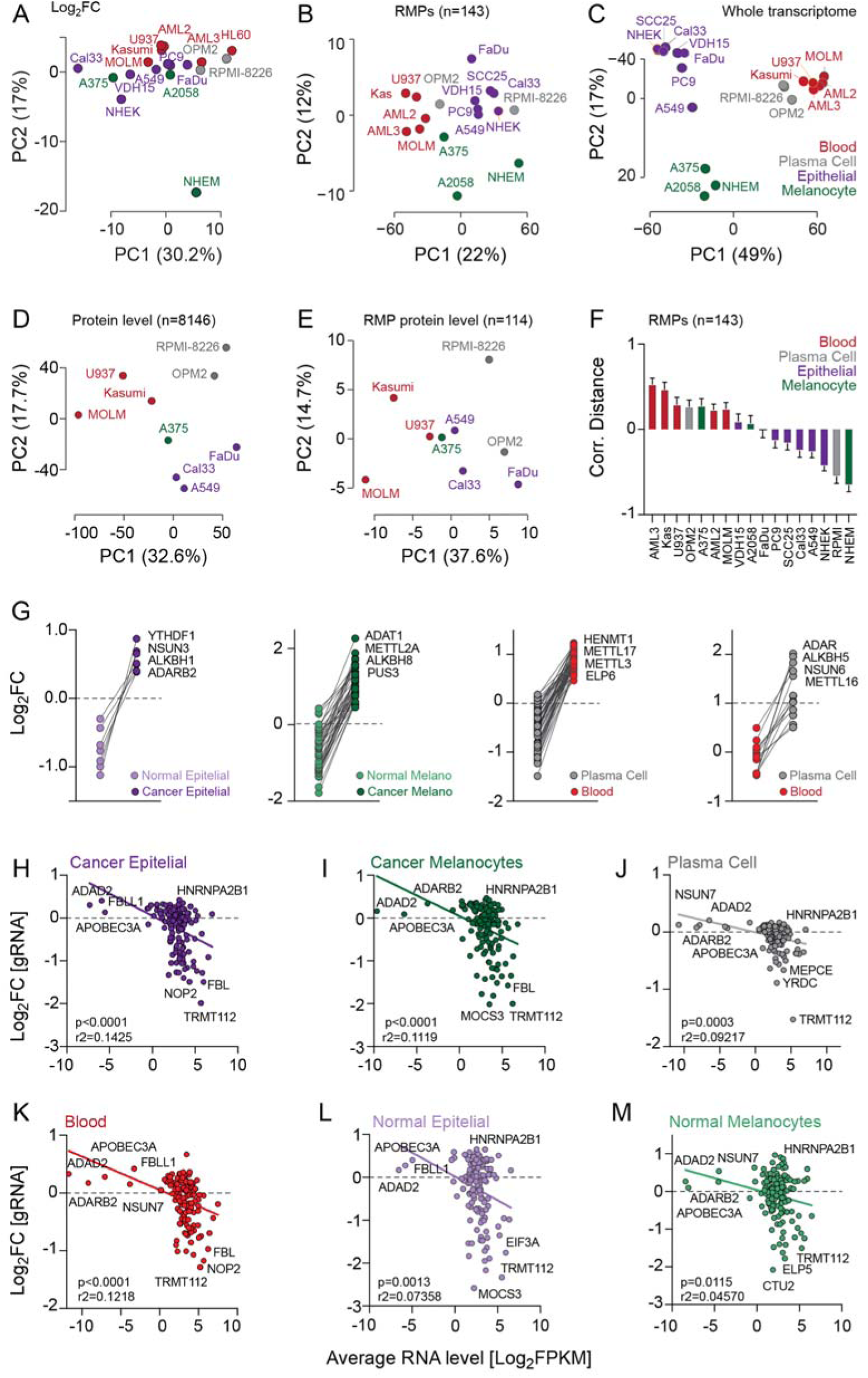
Essential RNA modifying proteins are not generally highly expressed. (**A**) Principal component analysis (PCA) using log_2_ fold change (FC) of 159 gRNAs in 18 cell lines originating from blood, plasma cells, epithelial cells or skin melanocytes at day 22 of the experiment showing cell line identify. (**B,C**) PCA using RMP expression (n=143) (B) or whole transcriptomes (C) (normalized log_2_ FPKM) in 18 cell lines (n=3 sequencing reactions per cell line) showing cell line identity. (**D,E**) PCA using expression levels of the whole proteome (n=81146) (D) or RMP protein expression (n=114) in the indicated cell lines (cBioPortal). (**F**) Correlation distance of RMP (n=143) RNA expression in the indicated cell lines. (**G**) Log_2_ fold change (FC) of tissue-specifically expressed RMPs in cancer cells originating from epithelia (purple), melanocytes (green), blood (red) or plasma cells (grey) compared to healthy controls (epithelia, melanocytes) or compared between blood cells (blood, plasma cells). (H-M) Correlation between log2 fold change (FC) gRNAs and RNA levels of 143 RMPs in cancer cells originating from epithelia (H), skin melanocytes (I), plasma cells (J), blood (K) or normal epithelial cells (L) and normal melanocytes (M).

**Supplementary Figure 4.**
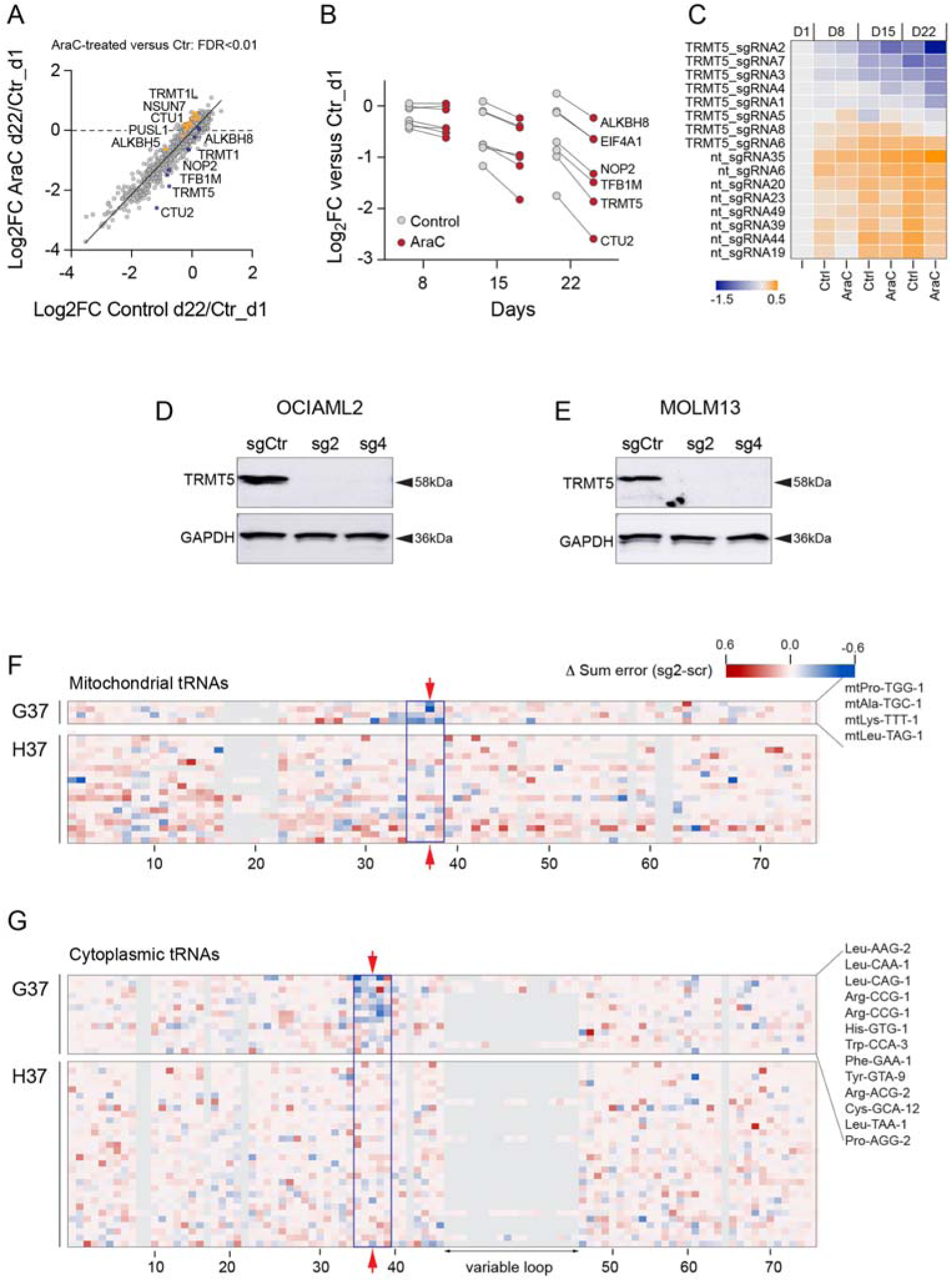
Synthetic lethality screen identifies novel function of TRMT5 in drug resistance. **(A)** Correlation of log2 fold change (FC) of sgRNAs in AraC-treated OCI-AML2 cells versus untreated cells at day 22 compared to day 1 of the experiment. Orange and blue dots: significantly (FDR<0.01) changes in abundance. (**B**) Log_2_ FC of sgRNAs targeting the indicated RNA modifying proteins (RMPs) in control (grey) and AraC-treated (red) OCI-AML2 cells after 8, 15, and 22 days of culture compared to day 1 of the experiment. (**C**) Heatmap showing log2 FC of all eight TRMT5-targeting and non-targeting (nt) sgRNAs at the indicated experimental days. (**D,E**) Western blots showing TRMT5 protein expression levels in OCI-AML2 (D) and MOLM13 (E) control (sgCtr) and knockout (sg2 and sg4) cells. GADPH serves as loading control. (**F,G**) Heatmap of summed error of read counts in mitochondrial (F) and cytosolic (G) tRNAs with (G37) or without (H37) a guanosine at position 37 in TRMT5-knockout (sg2) over control (scr) OCI-AML2 cells.

**Supplementary Figure 5.**
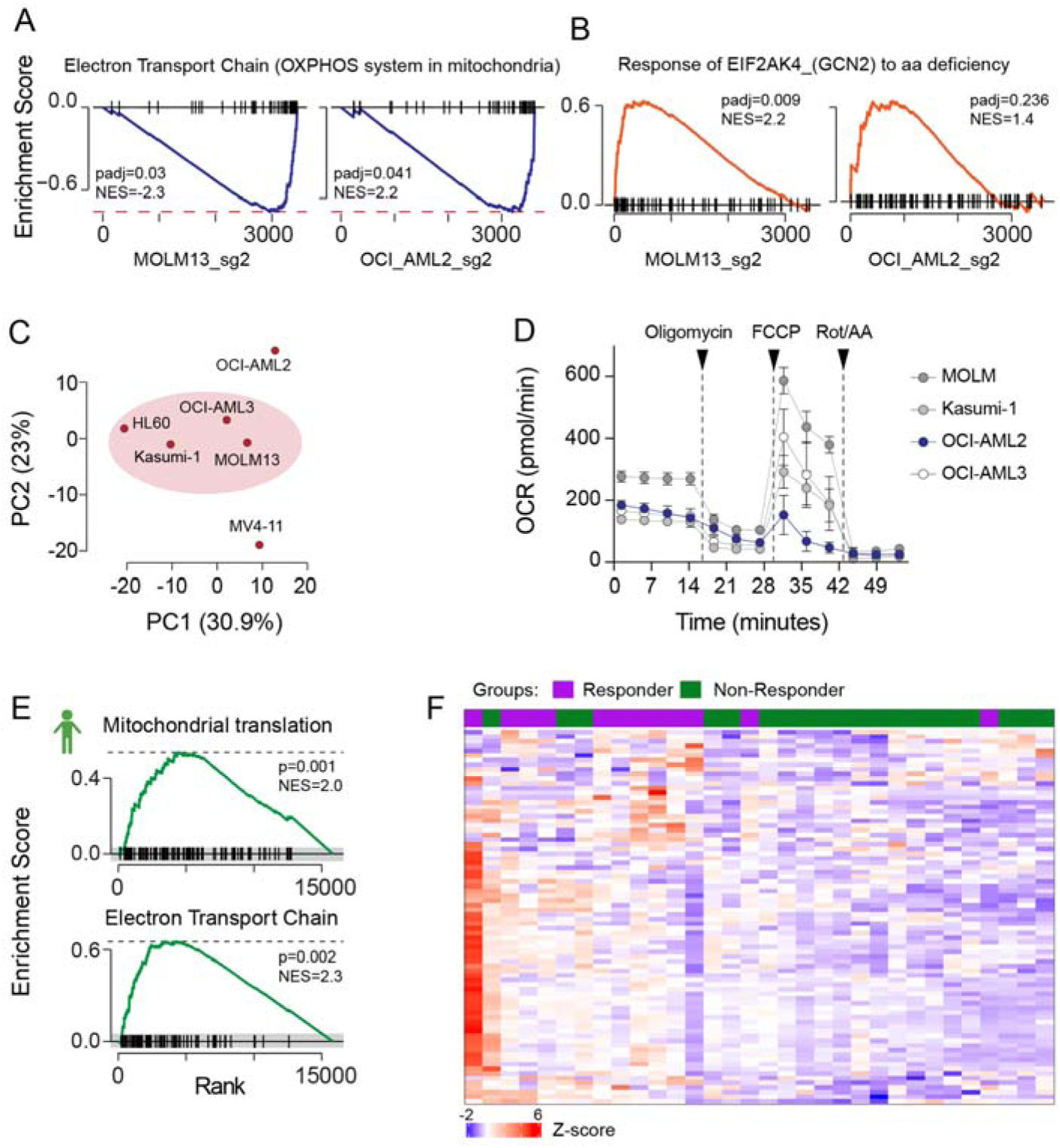
Low mitochondrial energy production in TRMT5-knockout and drug-tolerant AML cells. **(A,B)** Gene Set Enrichment Analysis (GSEA) of underrepresented proteins of the electron transport chain (A) and overrepresented proteins of the amino acid (aa) starvation stress response (B) in OCI-AML2 and MOLM13 TRMT5-knockout cells (sg2) compared to control cells infected with a scramble guide RNA. (**C**) Cluster analyses using mitochondria protein levels (MitoCarta3.0) indicated cell lines. (**D**) Oxygen consumption rates (OCR) in the indicated cell lines. (**E**) GSEA analyses for genes encoding protein regulating mitochondrial translation (top) and the electron transport chain (bottom) using RNA levels of AraC-treatment responsive leukemia patients at the time of diagnosis (n=12) showing complete remission after first induction therapy. (**F**) Heatmap showing z-score of RNA expression levels of genes encoding or the citruc acid (TCA) cycle and respiratory electron transport (n=80) in responding (n=12) (purple) and non-responding (n=20) (green) patients after first induction therapy.

**Supplementary Figure 6.**
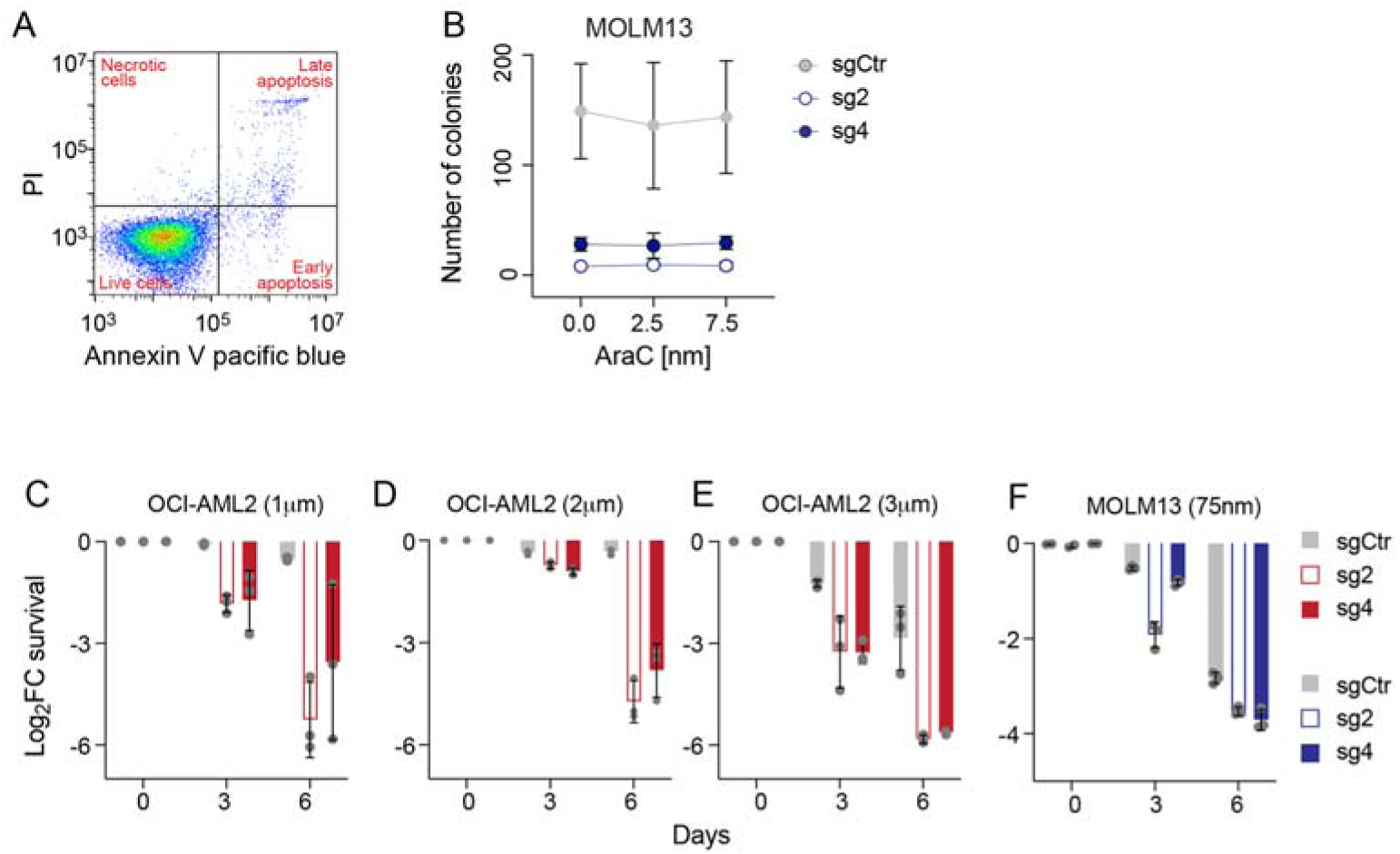
TRMT5-knockout re-sensitizes to drug treatment. **(A)** Representative scatter blot showing gating strategy to measure apoptosis using Annexin V labelling. PI: Propidium iodide. **(B)** Colony-forming efficiency (CFE) assays of MOLM13 control (sgCtr) and knockout (sg2, sg4) cells in the presence of increasing concentrations of AraC. (**C-F**). Second independent experiment showing log_2_ fold change (FC) of survival of drug-tolerant OCI-AML2 (I-K) and drug-sensitive MOLM13 (L) control (sgCtr) and TRMT5-knockout (sg2, sg4) cells at the indicated time points.

